# Using physiologically-based models to predict *in vivo* skeletal muscle energetics

**DOI:** 10.1101/2024.05.21.595083

**Authors:** Ryan N. Konno, Glen A. Lichtwark, Taylor J. M. Dick

## Abstract

Understanding how muscles use energy is essential for elucidating the role of skeletal muscle in animal locomotion. Yet, experimental measures of *in vivo* muscle energetics are challenging to obtain, so physiologically-based muscle models are often used to estimate energy use. These predictions of individual muscle energy expenditure are not often compared to indirect whole body measures of energetic cost. Here, we examined and illustrated the capability of physiologically-based muscle models to predict *in vivo* measures of energy use. To improve model predictions and ensure a physiological basis for model parameters, we refined our model to include data from isolated muscle experiments. Simulations were performed to capture three different experimental protocols, which involved varying contraction frequency, duty cycle, and muscle fascicle length. Our results demonstrated that these models are able capture the general features of whole body energetics across contractile conditions, but tended to under predict the magnitude of energetic cost. Our analysis revealed that when predicting *in vivo* energetic rates across contractile conditions, the model was most sensitive to the force-velocity parameters and the data informing the energetic rates when predicting *in vivo* energetic rates across contractile conditions. This work highlights it is the mechanics of skeletal muscle contraction that govern muscle energy use.

## 1 Introduction

Knowledge of how muscles consume energy to develop forces and perform mechanical work is fundamental in understanding the mechanisms that govern neuromuscular function and locomotor performance. This requires not only knowledge of the physiological processes that govern muscle energetics at the level of the fibre [Barclay and Curtin, 2023], but also the structure of the muscle and connective tissue [Curtin and Barclay, 2023], interaction with the external environment [Collins et al., 2015, Lichtwark and Wilson, 2007], and the neural control strategies used to perform a movement [Lai et al., 2018, Falisse et al., 2019]. An improved capacity to predict energetics during the diverse movements that humans and animals perform, and whether energy is optimized under certain conditions would afford insights into the mechanisms behind muscle energetic cost in a variety of locomotor tasks.

The energy used by skeletal muscle to perform mechanical work comes from chemical potential energy stored in ATP molecules; however, inefficiencies occur in the hydrolysis and rephosphorylization of ATP that result in the production of heat [Barclay, 2019]. To obtain measures of the heat released from muscle during contractions, experiments are typically performed on isolated animal muscle, whereby the heat can be directly measured using a thermopile (eg. Woledge et al. [1985], Barclay [1996]). While these experiments have provided us with invaluable information on the mechanisms governing muscle energetics at the muscle fibre level, these data are not easily extrapolated to *in vivo* human muscle. This is because the contractile conditions observed *in vivo* are often at submaximal activation levels and dynamic in nature, with time varying muscle fibre lengths and shortening rates; conditions very different to the controlled contractile environment of isolated muscle experiments. Our understanding of *in vivo* muscle energetics can be advanced using mathematical models in combination with experimental data [Umberger and Rubenson, 2011]. Modelling approaches are advantageous as they allow us to investigate direct cause and effect mechanisms, which is not often possible through experimental techniques.

A variety of approaches have been used to model the energetics of skeletal muscle during contraction. At the microscopic, single sarcomere level, Huxley-based models [Huxley, 1957] capture the dynamics of individual actin-myosin cross-bridges. While providing a detailed biophysical model, they are challenging to implement at larger scales owing to high computational costs and limited available experimental data that can be used to inform model parameters. This prompts the use of more phenomenological models, which are more appealing at a larger scale. At the whole body level, models have been developed based on measures of energetics from indirect calorimetry using O_2_ and CO_2_ measurements (e.g. Kram and Taylor [1990], Cavagna et al. [1977]). While these models are able to sufficiently capture whole body energetics, their highly phenomenological nature limits their potential to investigate the physiological principles that underlie muscle energetics and they are not transferable to predict movements beyond the original experimental protocol. Alternatively, physiologically-based, but still phenomenological, models (e.g. Lichtwark and Wilson [2005], Bhargava et al. [2004], Umberger et al. [2003]) serve as a middle ground utilizing rich datasets from single muscle experiments where fibre lengths and heat rates are directly measured. Within these models, the energy rates owing to maintenance, activation, and shortening/lengthening heats, as well as work are included (for more detail see Barclay and Curtin [2023]). These models have been applied to whole body musculoskeletal simulations of movement, such as walking [Umberger, 2010, Miller, 2014, Koelewijn et al., 2019] and hopping [Farris et al., 2014], but there can be large differences between model predictions and experimental measures of energy use [Miller, 2014, Koelewijn et al., 2019]; for example, during walking the predicted cost of transport varied from 2.45 *J* (*mkg*)^*−*1^ in the Bhargava et al. [2004] model to 7.15 *J* (*mkg*)^*−*1^ in the Lichtwark and Wilson [2007] model, whereas the measured cost was 3.35 *J* (*mkg*)^*−*1^ [Miller, 2014]. These differences could be owing to inaccuracies in model predictions of either energetics or forces, which remain difficult to predict during dynamic tasks such as walking. Under these conditions, it is particularly challenging to accurately model the high shortening rates and cyclical contractions, which are known to alter muscle energetics [Wakeling et al., 2023]. Thus, it could be more informative to predict energy use in simplified single joint tasks.

The purpose of this study was to investigate the ability of physiologically-based muscle models to predict whole body energy consumption from indirect measures of *in vivo* human muscle energetics. While similar validation studies have been performed previously for concentric and eccentric contractions [Umberger et al., 2003, Lentz-Nielsen et al., 2023], this study focuses on isometric conditions and identifying the mechanical mechanisms governing muscle energetic rates. To do this, we adapted and compared the model from Lichtwark and Wilson [2005, 2007] to a series of human experiments that investigated single joint contractions *in vivo* [van der Zee and Kuo, 2021, Beck et al., 2020, 2022], where the energy use was measured using indirect calorimetry. The existing model parameters were refined to represent physiological data obtained via direct measurements of energy use in isolated fibre-bundles from Barclay [1996]. By capturing the energetics from experiments *in vivo*, we aimed to identify if a physiologically-based model of skeletal muscle energetics could explain the increased energetic cost associated with changes in the rate of force development [van der Zee and Kuo, 2021], duty cycle [Beck et al., 2020], and muscle fascicle length [Beck et al., 2022]. Further, we investigated the sensitivity of our model to the input parameters to understand how the quality of experimental data impacts model predictions.

## 2 Materials and methods

Our mathematical model consists of a mechanical Hill-type model (subsection 2.1) in combination with the energetics model based on Lichtwark and Wilson [2005, 2007] (subsection 2.2). The mechanical component of the model was used to predict the muscle mechanics based on the forces measured within the experiment, and then the energetics component used the mechanical data to predict the whole muscle energetic cost (Figure 1). The model parameters were refined to capture physiological data from Barclay [1996] (subsection 2.3). These models were implemented in Python and are available at https://github.com/ryankonno/KLD2024-EnergeticsModel.

**Figure 1:**
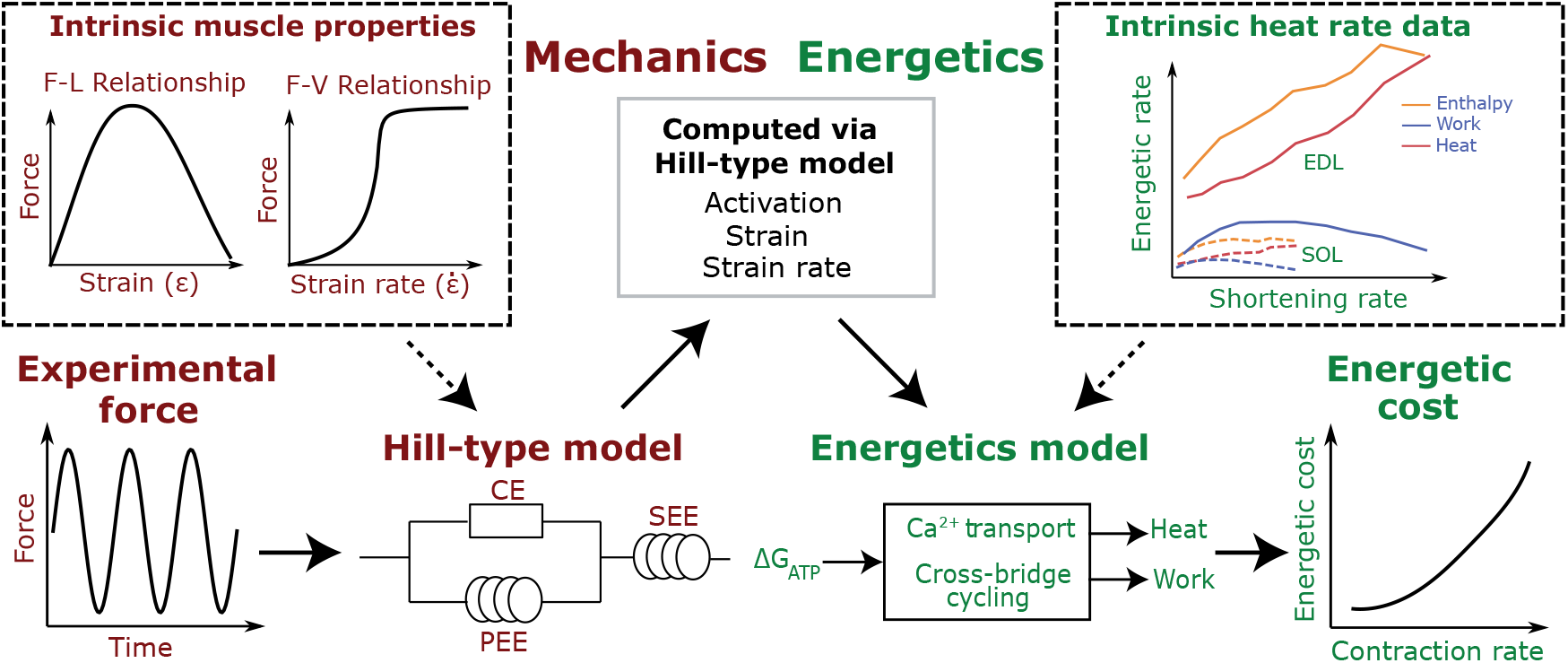
Overview of the modelling approach. For the experiments investigated in this study [van der Zee and Kuo, 2021, Beck et al., 2020, 2022], we used functions based on the experimentally measured forces as input to the Hill-type model. We then utilized a one-dimensional Hill-type model to solve for activation, strain, and strain rate. These variables were used as inputs to an energetics model, which is based on Lichtwark and Wilson [2005, 2007]. The heat rates are based on experimental data from Barclay [1996] in single muscle preparations of slow soleus (SOL) and fast extensor digitorum longus (EDL) muscles. The energetic cost of muscle contraction was then computed to determine the effect of the given experimental conditions: changes to contraction frequency [van der Zee and Kuo, 2021], duty cycle [Beck et al., 2020], or muscle fascicle length [Beck et al., 2022].

### 2.1 Mechanical model

We utilized a Hill-type muscle model that consists of a contractile element along with an in series elastic element, such that the total force from the muscle-tendon unit is given by

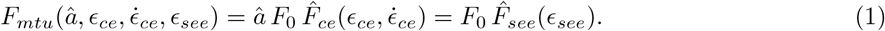

Here *F*_*mtu*_ is the total force from the muscle-tendon unit, *â* is the normalized activation in the muscle, and *F*_0_ is the maximum isometric force. The normalized force from the contractile element, 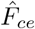, and series elastic element, 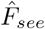, are written as a function of strains (*ϵ*_*ce*_ and *ϵ*_*see*_) and the strain rate 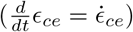. We neglected the presence of a parallel elastic element commonly seen in Hill-type models, as the experiments were all performed at isometric whole muscle length, whereby the muscle did not lengthen (*ϵ*_*ce*_ *≤* 0), and thus no contributions from the parallel elastic element (see e.g. Zajac [1989]). The function 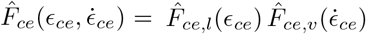 defines the intrinsic force-length and force-velocity properties of the muscle model, and uses the formulation from Dick et al. [2017].

### 2.2 Energetics model

We formulate the total energy rate of the muscle in terms of the muscle heat rate and work rate [Lichtwark and Wilson, 2005, 2007]. We used this model as our basis, but with the modification of the maintenance and shortening heat rate constants. These changes were made to obtain an energetics model that is entirely based on physiologically measured parameters. Here, the rates are normalized by *F*_0_ *l*_0_, where *l*_0_ is the resting fascicle length, to give normalized units of *s*^*−*1^. The total energy rate from the muscle is given as

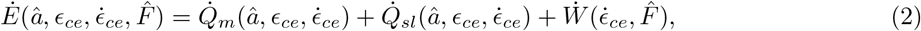

where 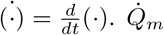 is the maintenance heat rate, 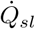 is the shortening and lengthening heat rate, and 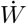 is the work rate. Here we use the term *maintenance heat* for what is typically referred to as the combination of heat from activation and generation of isometric force from the cross-bridges (see Barclay and Curtin [2023] for a review). 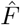 is the muscle force normalized to maximum isometric force, *F*_0_.

The maintenance heat rate is given by

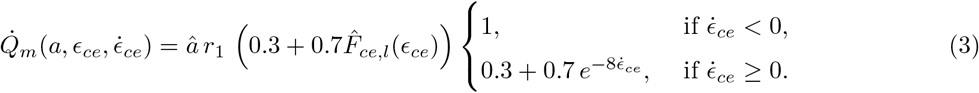

*r*_1_ is a parameter determined by experimental data from Barclay [1996] with units of *s*^*−*1^. This function represents the heat associated with Ca^2+^ flow and cross-bridge cycling under isometric conditions, but is scaled to account for decreased heat rates in lengthening muscle [Linari et al., 2003]. In addition to the isometric heat rates, there are typically increased heat rates associated with shortening and lengthening muscle [Hill, 1938, Barclay, 1996, Linari et al., 2003], which are given by

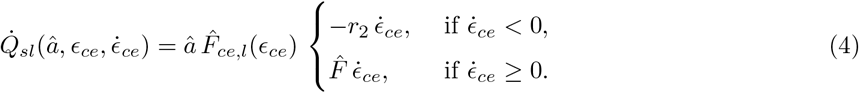

*r*_2_ is the unitless shortening heat rate parameter. In lengthening muscle there is no additional heat from cross-bridge cycling as is seen in shortening [Barclay and Curtin, 2023]; however, while the muscle lengthens, there is negative work being done on the muscle. All of this work is not transferred into elastic potential energy as the process is not completely efficient. Our model assumes that all work done on the muscle is eventually released as heat, as has been shown in experiments [Linari et al., 2003, Barclay and Curtin, 2023]. In addition to the heat rates, there is also the rate of work done by the contractile element, which is given by

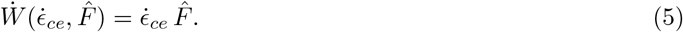

A summary of the parameters and variables used in the model, along with their sources are given in Table 1. All of the constants used in the model are based on experimental physiological data and are not optimised. The heat rates within the model are based on the intitial reactions and neglect the modelling of recovery heat rates. While these heat rates are smaller than the initial heats, they would increase the overall energetic rates [Barclay et al., 1995].

**Table 1:** Summary of model variables and parameters with definitions and units. Parameters used in this study were based on physiological data and not optimised.

| Variable | Definition | Source |
| --- | --- | --- |
| $l_0$ | Resting length of muscle ($m$) | Varied depending on experiment (see section 3) |
| $F_0$ | Maximum force at $l_0$ ($N$) | Calculated based on PCSA from Handsfield et al. [2014] |
| $\dot{\epsilon}_{ce,max}$ | Maximum strain rate ($s^{-1}$) | Estimated from experimental values Dick et al. [2017] |
| $\kappa$ | Force-velocity curvature | Slow-fibre values from Dick et al. [2017] |
| $\epsilon_{ce}$ | Strain of the CE | Calculated based on experiments (see section 3) |
| $\dot{\epsilon}_{ce}$ | Strain rate of the CE ($s^{-1}$) | Calculated from $\epsilon_{ce}$ |
| $\epsilon_{see}$ | Strain of the SEE | Solved for in the Hill-type model |
| $\hat{F}$ | Total muscle force (normalized by $F_0$) | Calculated using Equation 1 |
| $\hat{F}_{ce}$ | CE force (normalized by $F_0$) | Calculated using properties from Dick et al. [2017] |
| $\hat{F}_{see}$ | SEE force (normalized by $F_0$) | Calculated depending on the experiment (see section 3) |
| $k$ | Tendon stiffness ($N m^{-1}$) | Calculated depending on the experiment (see section 3) |
| $\hat{a}$ | Normalized activation | Calculated using Equation 1 |
| $\dot{E}$ | Total energy rate of the muscle ($s^{-1}$) | Calculated using Equation 2 |
| $t$ | Time ($s$) | Simulation length is based on experiment (see section 3) |
| $\dot{Q}_m$ | Maintenance heat rate ($s^{-1}$) | Calculated using Equation 3 |
| $\dot{Q}_{sl}$ | Shortening/lengthening heat rate ($s^{-1}$) | Calculated using Equation 4 |
| $\dot{W}$ | Work rate ($s^{-1}$) | Calculated using Equation 5 |
| $r_1$ | $\dot{Q}_m$ parameter ($s^{-1}$) | Optimised from Barclay [1996] (see Equation 3) |
| $r_2$ | $\dot{Q}_{sl}$ parameter ($s^{-1}$) | Optimised from Barclay [1996] (see Equation 4) |
| $s_f$ | Shift in muscle fibre length ($m$) | Defined based on data from Beck et al. [2022] |

### 2.3 Model parameters

The behaviour of the above model depends on *r*_1_ and *r*_2_, which are the constants in the maintenance and shortening heat rates, respectively. To determine these parameters, we used existing data on the thermodynamic behaviour of muscle during shortening contractions. Barclay [1996] measured both muscle work and heat rates in single muscles consisting mainly of either slow- or fast-type fibres. The slow fibre-type data are from experiments on mouse soleus (SOL), while the fast fibre data are from mouse extensor digitorum longus (EDL). To determine our model parameters, we minimized the relative error between predicted model heat rates and the experimental data (see Supplementary Material for more detail). We first scaled the experimental data to account for temperature differences. The experiment was conducted at 25^*°*^C, whereas *in vivo* temperatures are near 35^*°*^C. This required us to increase the heat rates by a factor of 4 [Rall and Woledge, 1990] and the maximum strain rate *ϵ*?_*ce,max*_ by a factor of 2 [Ranatunga, 1998]. The resulting parameters are shown in Table 2.

**Table 2:** Values for the energetic parameters, *r*_1_ (*s*^*−*1^) and *r*_2_ (unitless). The parameters were obtained via fit to physiological data from Barclay [1996]. *r*_1_ and *r*_2_ are the maintenance heat and shortening heat parameters, respectively.

| | $r_1$ | $r_2$ |
| --- | --- | --- |
| Slow-type muscle (SOL) | 0.618 | 0.234 |
| Fast-type muscle (EDL) | 2.792 | 0.697 |

### 2.4 Computational experiments

To test the model, we compared predictions of muscle energy consumption to whole body energy consumption obtained via indirect calorimetry. We set up our model to capture three different experimental paradigms that investigated the influence of (i) rate of force development [van der Zee and Kuo, 2021], (ii) duty cycle [Beck et al., 2020], and (iii) muscle fascicle length [Beck et al., 2022] on muscle energetics. The first study, van der Zee and Kuo [2021], demonstrated that the energetic cost increased with faster cycle frequencies during a seated knee extension task. Beck et al. [2020] examined the effect of duty cycle, the fraction of the contraction cycle where the muscle is actively producing force, on energetics during plantarflexion contractions with knee flexed to 50^*°*^ to isolate the contribution from the SOL. They found that energetic cost increased as duty cycle decreased. Beck et al. [2022] explored the effect of fascicle length, owing to differences in ankle angle during cyclical isometric plantarflexion contractions, on energetic cost. They found that an increased ankle angle corresponded to shorter fascicle lengths, which led to increases in energy consumption. These experimental studies were chose to evaluate our model predictions because they elicit different energetic responses and each measured relevant parameters that influence muscle energy use, namely muscle activation level (via EMG) and fascicle length changes of at least one of the muscles contracting (via ultrasound imaging). We will refer to the three simulations as vdZ2021, B2020, and B2022 for the respective experimental conditions. The specific model parameter values used for each of these experiments are provided in the Supplementary Material.

#### 2.4.1 van der Zee and Kuo 2021

van der Zee and Kuo [2021] investigated the effect of contraction frequency on the energetic cost of isometric muscle contraction. In their experiment, participants were seated and performed bilateral knee isometric extensions following a sinusoidal torque trace, at frequencies ranging from 0.5 *Hz* to 2.5 *Hz* (Figure 2A). The main driver behind whole body energetic cost is assumed to be the knee extensor muscles. Thus, we implemented our model as a lumped muscle model for the combined vastus lateralis, vastus intermedius, vastus medialis, and rectus femoris; for one lumped muscle, we would expect it to match half of the total energetic cost reported. We used the *l*_0_ value from the vastus lateralis of *l*_0_ = 0.095 *m* [van der Zee and Kuo, 2021] for the lumped muscle, since strains and activation levels are reported in van der Zee and Kuo [2021] for this muscle, and we use *F*_0_ = 4775 *N* based on physiological cross-sectional area (PCSA) calculated in Handsfield et al. [2014]. The tendon stiffness of the model (*k* = 1.58 *×* 10^4^ *N m*^*−*1^) was prescribed to achieve experimentally measured fascicle strains over a cycle (*ϵ*_*max*_ = *−*0.147). Starting with a sinusoidal torque matching the experimental conditions, we solved the Hill-type model for the time-varying strains, strain rates, and activation levels (Figure 1). These data were then used by the energetics model to predict the energetic cost.

**Figure 2:**
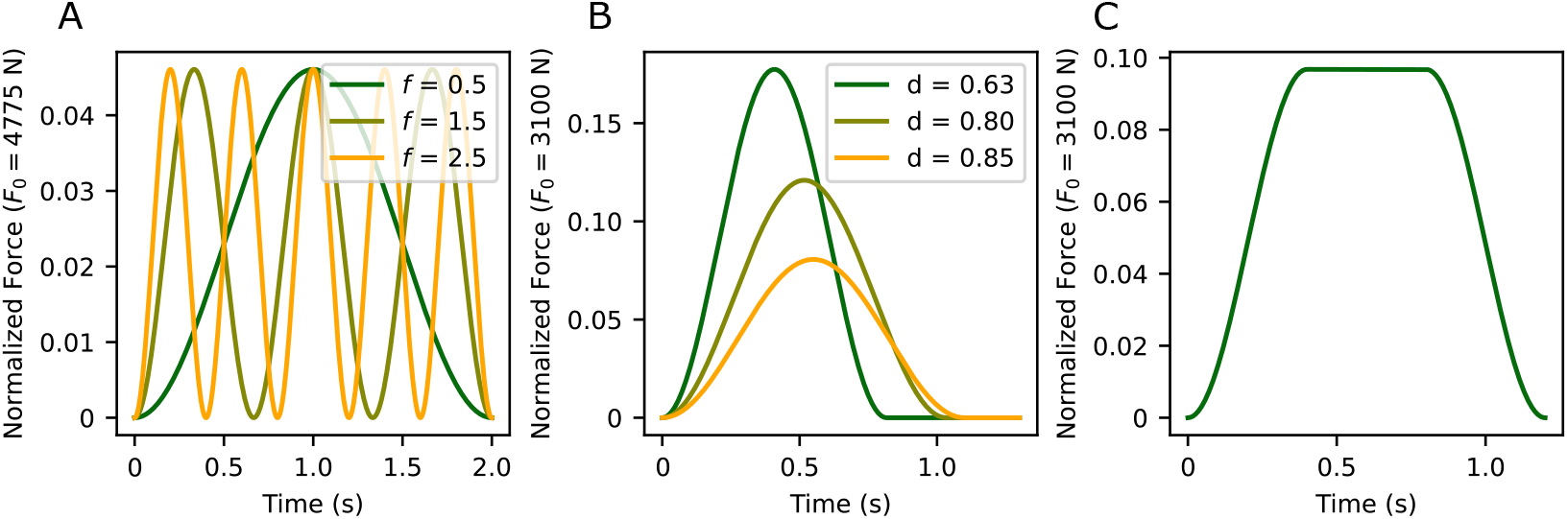
Experimentally based force used as input to the Hill-type model. vdZ2021 varies the frequency of contraction (A). B2020 investigate the effect of duty cycle, while maintaining the same integral of the force with respect to time by increasing the peak force (B). B2022 simulations maintain the same force input while decreasing muscle fascicle lengths (C).

#### 2.4.2 Beck et al. 2020

The Beck et al. [2020] experiments investigated the influence of duty cycle on energetic cost. Here we used the duty cycles reported in the experiment of 0.63, 0.8, and 0.85 [Beck et al., 2020] (Figure 2B). We provided our model with a force trace based the the predicted SOL force in [Beck et al., 2020] Figure 3. Our model used an *l*_0_ = 0.0386 *m* based on data provided in Beck et al. [2020] and *F*_0_ = 3100 *N* similarly calculated using PCSA values from Handsfield et al. [2014]. Here we aimed to account for the energy cost owing only to the SOL muscle, so we scaled the input force and the experimental energetic cost. This was done by assuming that force contributions to the total torque from each of the muscles was based on the relative PCSA of the triceps surae muscles: the medial (MG) and lateral (LG) gastronemii and the SOL. While Beck et al. [2020] suggested that the majority of the energetic cost is due to the SOL, they reported large activation levels in the LG (e.g., low duty high torque condition: peak LG activation *≈* 35 *mV*, peak SOL activation *≈* 45 *mV* [Beck et al., 2020], and did not measure EMG in the MG). Because of this, we made the assumption that the the LG and MG have equal contributions to the energetic cost, which is supported with experimental EMG data from one participant (see Supplementary Material). Using PCSA values from Handsfield et al. [2014] (124 *cm*^2^ (SOL), 50 *cm*^2^ (MG), and 23 *cm*^2^ (LG)), we can scale the total energy by the SOL contribution to the total PCSA (60%). The width of the force-length relationship from Dick et al. [2017] was decreased to achieve a better comparison to the Beck et al. [2022] results; this was done for both the B2020 and the B2022 simulations. Tendon stiffness (*k* = 15.5 *×* 10^4^ *N m*^*−*1^) was set to approximate the maximal experimental fascicle strain (*ϵ*_*max*_ *≈* 0.05 to 0.1).

**Figure 3:**
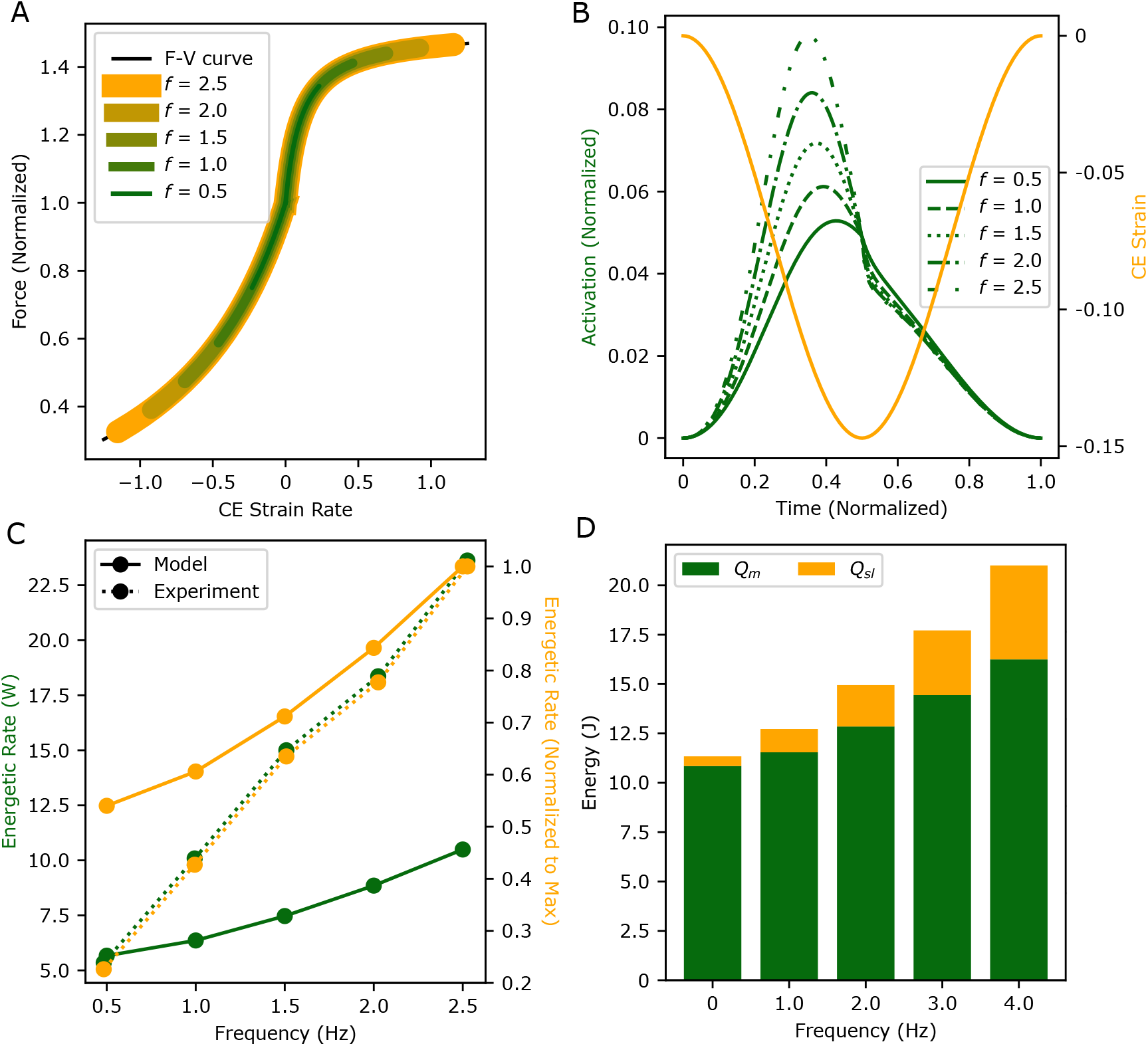
Modelling results for the vdZ2021 simulations investigating the effect of contraction frequency on energetic cost. The range of the force-velocity relationship traversed by the contractile element increases with increasing contraction cycle frequency (A). Muscle strains were constant between frequencies, but activation increased with frequency (B). Comparing the relative changes in energetic cost with cycle frequency, the model predicted *≈* 175% increase in the energetic cost, which nearly matched the experimental results (C, yellow lines). The absolute energetic cost values were lower than the experimental values (C, green lines). The relative contribution of the shortening heat (*Q*_*sl*_) to the total heat output increased with frequency (D).

#### 2.4.3 Beck et al. 2022

Beck et al. [2022] found an increase in energetic cost when muscle operates at shorter fascicle lengths. Similarly to the B2020 simulations, we scaled the input forces and experimental energetic cost. We used an input force based on the experimental condition provided in Beck et al. [2022] (see Figure 2C). Here, plantarflexion contractions were performed with the knee angle at 50^*°*^ to isolate contributions from the SOL [Beck et al., 2022]. In the experiments, a resting state (ankle angle = 90^*°*^) corresponded to a SOL fascicle length of 0.0415 *m*, whereas at greater ankle angles (ankle angle = 120^*°*^) the resting fascicle length was 0.035 *m*. We used a similar modelling approach as the Beck et al. [2020] experiment, but to model the changes in fascicle lengths observed in the experiment, a fascicle length shift was incorporated into the calculation of the contractile element strain,

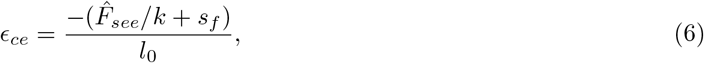

where *s*_*f*_ is the shift in the fascicle length (in *m*) and *k* is the normalized tendon stiffness (in *m*^*−*1^). To capture the three conditions we used fascicle shifts *s*_*f*_ of 0, 0.1 *l*_0_, and 0.2 *l*_0_. Tendon stiffness (*k* = 15.5 *×* 10^4^ *N m*^*−*1^) was set to maintain maximum strain *ϵ*_*max*_ *≈* 0.05 from the resting length for a given ankle angle.

## 3 Results

### 3.1 van der Zee and Kuo 2021

Our model predicted that as the frequency of contraction increased, the contractile element of the model traversed a larger region of the force-velocity relationship owing to the greater strain rates at higher contraction frequencies (Figure 3A). This resulted in a higher muscle activation required to maintain the same force over the contraction cycle (Figure 3B). Here we observed an increase in activation of 85% from 0.5 *Hz* to 2.5 *Hz*, which is similar to the 75% increase measured experimentally [van der Zee and Kuo, 2021]. van der Zee and Kuo [2021] found an increased energetic cost of 18.15 *W* (for one leg) when the contraction frequency increased from 0.5 *Hz* to 2.5 *Hz*. Despite a qualitatively similar trend to these results, our model predicted a smaller difference in energetic cost of 4.82 *W* between conditions (Figure 3C, dotted line). As the contraction frequency increased, the higher activation levels resulted in an increase in the maintenance heat, while the faster shortening rates resulted in larger contributions from the shortening heat (Figure 3D).

### 3.2 Beck et al. 2020

With a decrease in duty cycle, we observed an increase in the maximum strain rate over the contraction cycle (0.12 *s*^*−*1^ to 0.35 *s*^*−*1^), which corresponded to a larger region of the force-velocity relationship traversed in the model predictions (Figure 4A). The increased strain rates resulted in an increase in activation level (from *â ≈* 0.08 to *â ≈* 0.23) to maintain a given force (Figure 4B). Beck et al. [2020] did not report the normalized activation levels, but when comparing the relative change in activation, the model displayed a 150% increase in *â* and experimentally there was an increase of *≈* 75%. The peak contractile element strains over a cycle predicted by the model were 0.091, 0.063, and 0.042 for duty cycles of 0.63, 0.80, and 0.85, respectively, while the experimental peak muscle fibre strains were 0.098, 0.081, and 0.056. The model under-predicted the energetic cost from these experiments, as it only reached *≈* 10% of the scaled experimental values (Figure 4C); however, when normalized to the peak energetic rate, we found that there was a very similar qualitative trend in the energetic cost across duty cycles (Figure 4C). There were increases in both the maintenance and shortening heat contributions to the total energy use, but higher strain rates led to a larger proportion of the total energy contributed by the shortening heat (Figure 4D), consistent with the vdZ2021 simulations.

**Figure 4:**
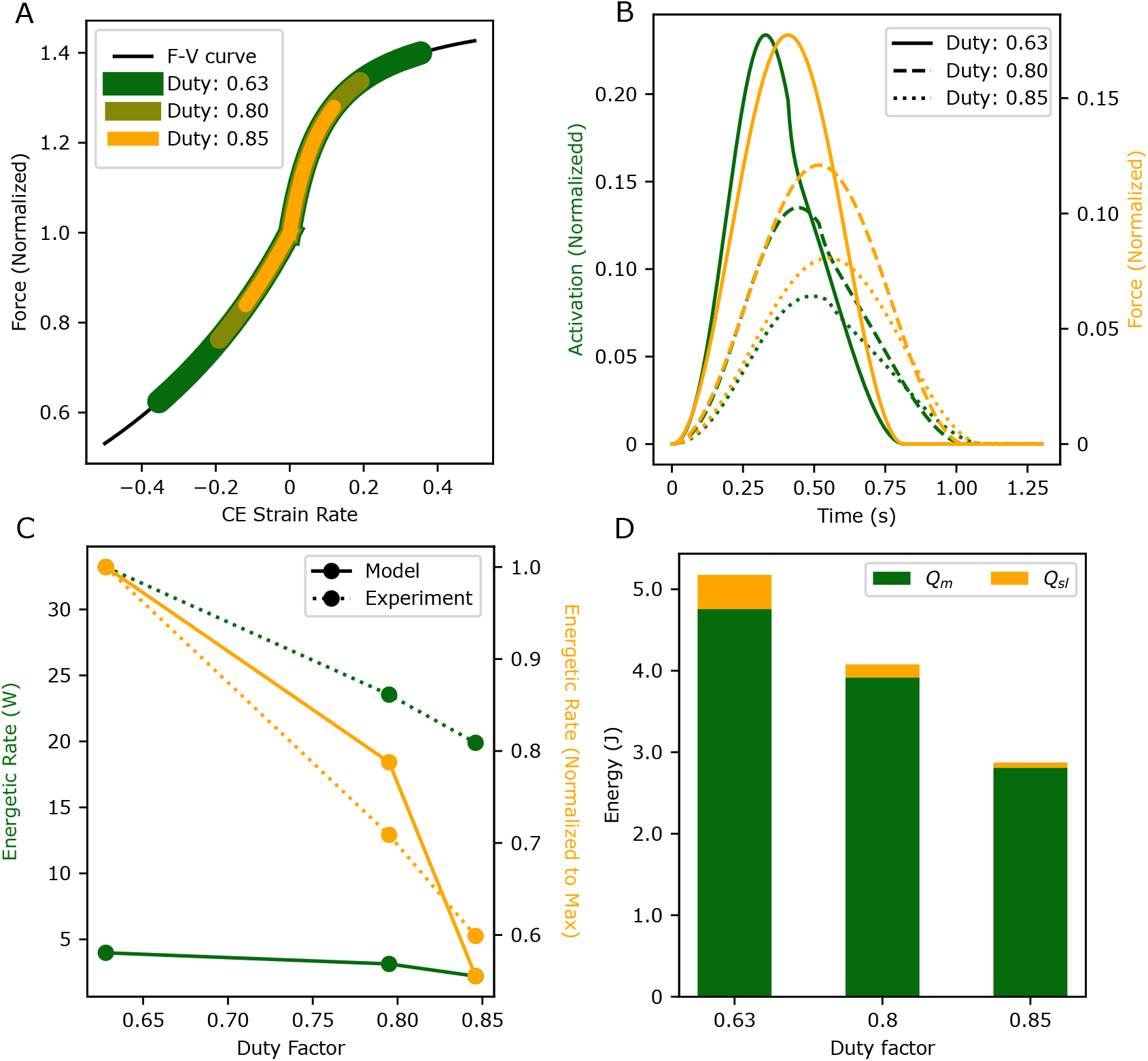
Results from the B2020 simulations investigating the effect of changes in duty cycle on energetic cost. The regions of the force-velocity relationship traversed and maximum activation increased with decreasing duty cycle (A and B, respectively). The energy rates showed an increase with decreasing duty factor, as shown by the relative changes in energetic rate (C, yellow lines). However, the magnitude of energetic cost did not match the experimental data from Beck et al. [2020] (C, green lines). There was a larger contribution from the shortening heat (*Q*_*sl*_) with lower duty cycles (D).

### 3.3 Beck et al. 2022

The B2022 simulations investigated the effect of muscle fascicle operating length on energetic cost, and found that with increasing contractile element strain (modelling decreased fascicle lengths) there was a leftward shift in operating range on the force-length relationship (Figure 5A), which led to large increases in muscle activation. To produce an equivalent peak force, the peak normalized activation increased from *â* = 0.105 to 0.309 when resting fibre strain factor was increased from *s*_*f*_ = 0 to 0.2 *l*_0_ (Figure 5B). This is approximately the same relative increase in activation levels, but smaller magnitudes than observed experimentally (0.25 to 0.77 increase in activation [Beck et al., 2022]). The peak fascicle strains predicted by the model were 0.047, 0.147, and 0.247 for *s*_*f*_ = 0 *mm*, 4 *mm*, and 8 *mm*, while the experimental peak fascicle strains were 0.05, 0.15, and 0.22 for initial shifts in fascicle length of 0 *mm*, 2.9 *mm*, and 6 *mm*, respectively [Beck et al., 2022]. The peak strain rates of 0.18 *s*^*−*1^ observed was the same between conditions, and *≈* 30% smaller than the experimental results (see Beck et al. [2022] Figure 5B). The energetic cost of these contractions showed a similar trend to the experimental data but was much smaller in magnitude. Shorter fascicle lengths resulted in an *≈* 50% increase in energetic rate, while experimental measures showed *≈* 200% increase (Figure 5C).

**Figure 5:**
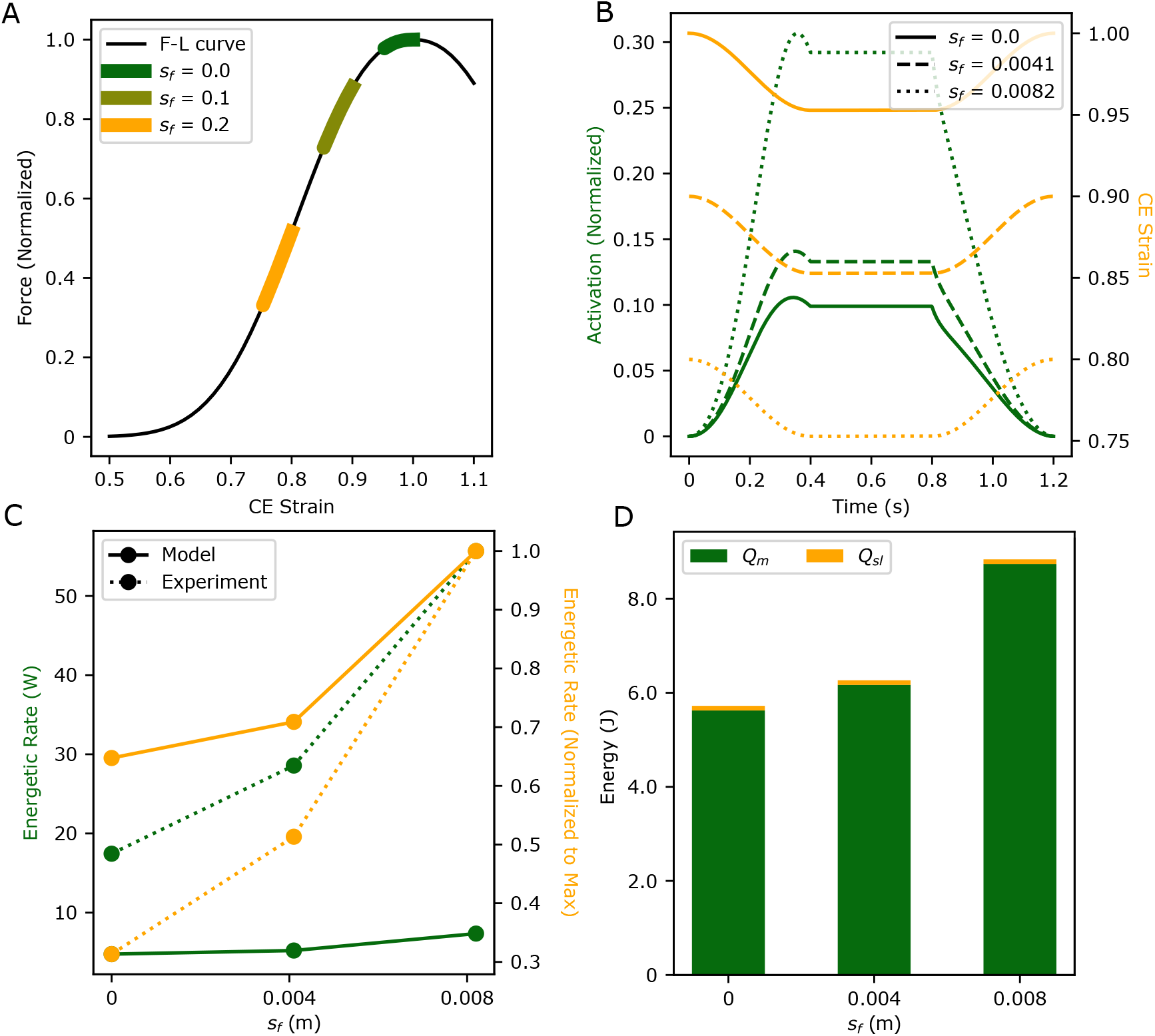
Modelling results from the B2022 simulations investigating the effect of changes to fascicle length on energetic cost. There was a shift in region of the force-length relationship occupied with a shift in the fascicle length (A). Activation levels increase with larger shifts in fascicle length (B). When comparing the relative change in the heat rates (C, yellow lines), there is a similar trend to the experimental data. The absolute values for the energetic rates predicted by the model have a lower magnitude (C, green lines). The relative contribution from the heat components (*Q*_*sl*_ and *Q*_*m*_) does not change between conditions (D).

The difference in activation level between conditions was largely responsible for alterations in energetic cost, as the shortening heat rate was unchanged under varying *s*_*f*_ (Figure 5D).

### 3.4 Sensitivity analysis

To understand the influence of specific modelling parameters on the muscle model energy predictions, we performed a sensitivity analysis (Figure 6). We chose to perform the sensitivity analysis on the parameters that demonstrate large experimental variation and are challenging to experimentally measure: the energetic parameters *r*_1_ and *r*_2_, and the force-velocity constants, *κ* and 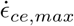. This analysis consists of (i) a relative sensitivity analysis (Figure 6Ai, Bi) during a shortening contraction whereby the muscle was activated to 100% activation and then allowed to shorten at fixed velocities of 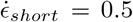, 1.0, and 1.5 *s*^*−*1^ over the plateau of the force-length relationship and (ii) a sensitivity analysis (Figure 6Aii, Bii) whereby we investigated the sensitivity of the predicted energy rate under the vdZ2021, B2020, and B2022 simulation protocols to the input parameters. The relative sensitivities correspond to a ratio between a percent change in the parameter and a percent change in the predicted energy use; in particular, 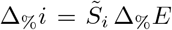, where Δ_%_*i* is the percent change in parameter *i*, 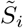 is the relative sensitivity to *i*, and Δ_%_*E* is the percent change in the energy.

**Figure 6:**
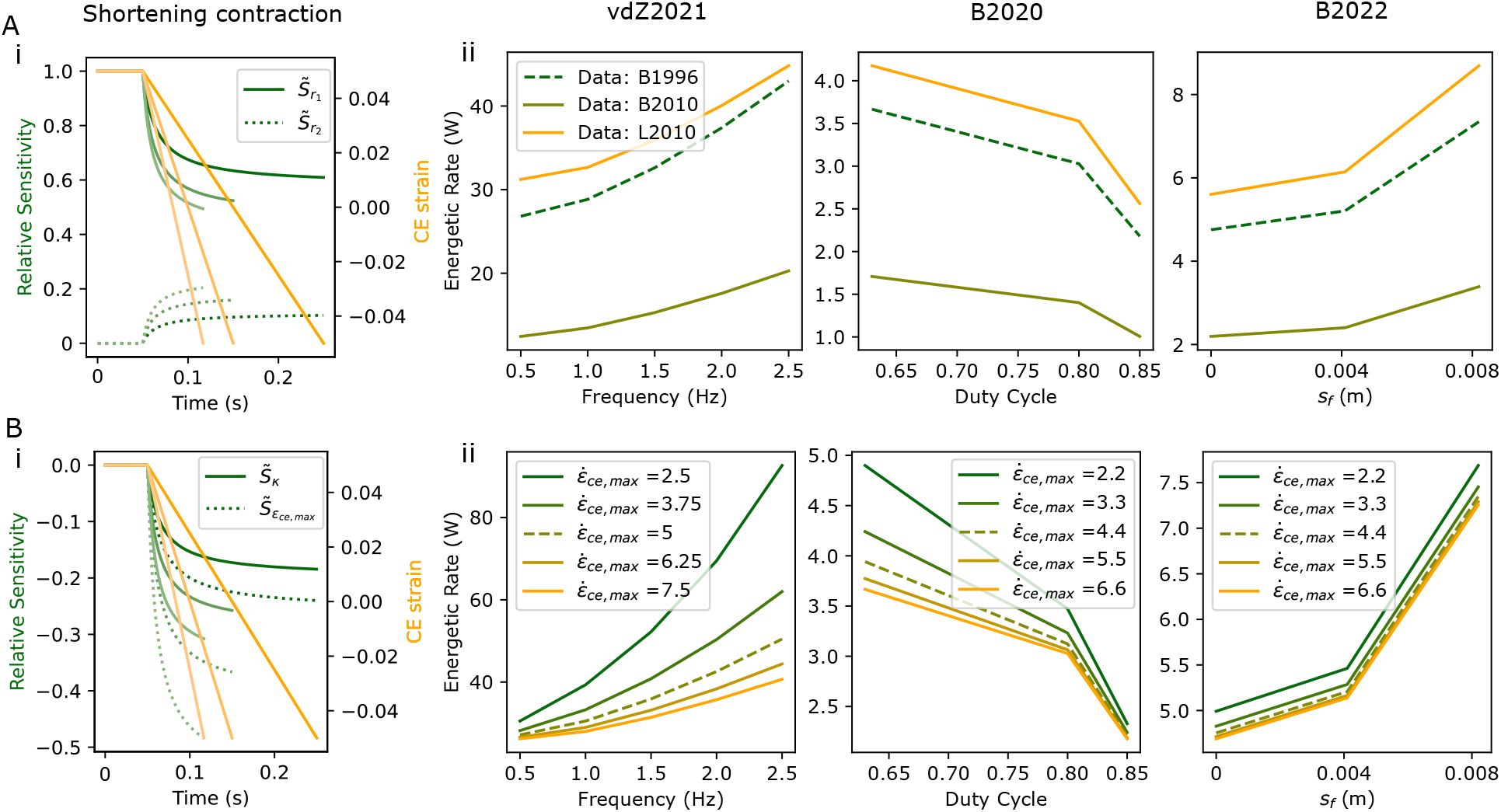
Model sensitivities during shortening contractions and the vdZ2021, B2020, and B2022 simulation protocols. Sensitivity to energetic parameters (A). The relative sensitivity to the energetics parameters, *r*_1_ (solid lines) and *r*_2_ (dotted lines), are shown in (Ai) and the sensitivity to the energetic data used to determine model parameters is shown in (Aii). These data were taken from Barclay [1996] (B1996), Barclay et al. [2010] (B2010), and Lichtwark and Barclay [2010] (L2010). Sensitivity in the force-velocity parameters (B). Relative model sensitivity to the force-velocity curvature, *κ* (solid lines), and the maximum strain rate, 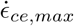 (dotted lines), are shown in (Bi). 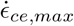 was varied by *±*25% and *±*50% in (Bii). Dashed lines represent the base parameter set used for previous experiments. The relative sensitivities were calculated as the ratio of a percent change in the parameter to a percent change in the predicted energy use. Here, we use 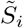 to denote the relative sensitivity with respect to parameter *i*. Shortening contractions were simulated at shortening rates of 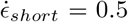, 1.0, 1.5 *s*^*−*1^, which corresponds to fading colour on the plots (dark to light), and using slow-type fibre properties.

The sensitivity analysis on the energetic parameters demonstrated that the model was most sensitive to the maintenance heat parameter, *r*_1_, during the isometric contraction with a relative sensitivity 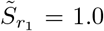 while 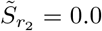 (Figure 6Ai). During the shortening period, energy use became less sensitive to *r*_1_ and the sensitivity to the shortening heat rate parameter, *r*_2_, increased, while still maintaining higher sensitivity to *r*_1_ (Figure 6Ai). Given the high sensitivity to these parameters, we examined how using different data to inform these parameters would influence the outputs in the vdZ2021, B2020, and B2022 simulation protocols. In addition to the Barclay [1996] parameter values given in Table 2, the energetics parameters were determined based on the data from Barclay et al. [2010] and Lichtwark and Barclay [2010]. This effectively corresponds to a range in *r*_1_ from 0.38 to 0.74*s*^*−*1^ and a range in *r*_2_ from 0.09 to 0.23, demonstrating high variability in the data used to derive energetic model parameters. Considering these datasets, we find that the predicted energetic rates can vary by *≈* 150% depending on the experimental condition (Figure 6Aii). The datasets considered here measured heat rates in slow twitch soleus muscle during fibre-bundle experiments (for the methods used to determine parameter values see the Supplementary Material).

The sensitivity analysis on the force-velocity relationship curvature, *κ*, and maximum shortening rate, 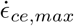 demonstrated that during shortening the model energy use was more sensitive to 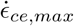 compared to *κ* (Figure 6Bi). Note that the negative relative sensitivity values indicate a percent change in the variable will result in the corresponding decrease in the energetic cost. As expected, during an isometric contraction (with no tendon), there was no influence of *κ* and 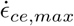 on energy use (Figure 6Bi). Given the model was more sensitive to 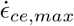, we examined the sensitivity during the vdZ2021, B2020, and B2022 simulations by varying 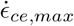 by *±*25 and *±*50%. Figure 6 (Bii) demonstrates the strong sensitivity on 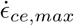 at the high contraction rates observed in vdZ2021 and B2020 protocols. B2020 simulations were less sensitive to 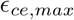 with no change in the magnitude of the sensitivity between conditions, which is expected based on limited changes to the shortening rates (see Figure 5).

## 4 Discussion

This study investigated the ability of a physiologically-based muscle model [Lichtwark and Wilson, 2005, 2007] to capture the energetics of *in vivo* skeletal muscle contraction under a range of mechanical conditions [van der Zee and Kuo, 2021, Beck et al., 2020, 2022]. Previously, these models have been shown to accurately predict energetics of isolated muscle, but have performed poorly when compared to *in vivo* energetic rates for single joint contractions [Lentz-Nielsen et al., 2023]. This warrants an investigation of the mechanisms governing the muscle energetic rates *in vivo*. In many cases, these models have been applied to study whole body energetics during complex locomotor tasks [Miller, 2014, Koelewijn et al., 2019]; however, during simple single joint movements, whereby the neuromechanical contributions of an individual muscle or single group of muscles can be interpreted, model predictions of *in vivo* energetics is not completely understood. Our results demonstrate the physiologically-based models are able to capture the general patterns in energetic rates measured experimentally, but the absolute values tend to vary considerably from measures of muscle energy use from indirect calorimetry. This experimentally-informed modelling framework provides rich insights into muscle energetics, but the discrepancies between model predictions and experimental measures warrants further exploration.

### 4.1 Energetic rate predictions

In the vdZ2021 simulations, the energetic rates calculated by the model were able to predict the trends in the experimental data, but the magnitude differed between the model predictions and the experiments. van der Zee and Kuo [2021] found that the model by Uchida et al. [2016] was only able to account for *≈* 7% of the change in energetic cost measured in their experiments, leading to the suggestion that other mechanisms are required to explain the energetic cost associated with the rate of force development. Here, we find that the Lichtwark and Wilson [2005, 2007] model with experimentally based parameters, while only predicting *≈* 25% of the experimentally measured energetic cost, does capture the trends in energetic cost observed with the increased contraction frequency. There are a number of possible reasons for the discrepancy in absolute energetic rate, which could range from assumptions about the sources of energy in the experiment (e.g. co-contraction in other muscles at fast cycle frequencies) to inadequate scaling of parameters to whole muscle energetics. Further, the model neglects the oxidative recovery heat rates, which are a result of ATP regeneration within the mitochondria [Barclay and Curtin, 2023]. As the experiments [van der Zee and Kuo, 2021, Beck et al., 2020, 2022] took place over a period of 5 minutes, it is likely that there would be contributions from the recovery heat, which may help to explain the difference between the measured experimental energy rates and model predictions.

The values of energetic rate measured in the single joint studies by Beck et al. [2020] and Beck et al. [2022] are potentially high considering the simplicity of the task (e.g. metabolic rate is 20% of net walking metabolic rate [Umberger, 2010]). There are a number of possible explanations, particularly relating to the high chance of metabolic energy consumption occurring due to other muscle contractions and/or physiological processes (e.g. breathing). For example, there is likely a high amount of co-contraction occurring in other lower limb muscles, such as the more proximal and larger hamstrings and quadriceps muscles, or in the postural muscles that stabilized the body. Beck et al. [2020] did show that there was a large amount of co-activation in the antagonist tibialis anterior (TA), which was not accounted for in our scaling of the experimental energetic rates. To verify that co-contraction was occurring in the other lower limb muscles, we conducted a pilot experiment in one participant (see the Supplementary Material), where we replicated the Beck et al. [2020] protocol, but with sEMG measurements on the ankle plantar flexors, ankle dorsiflexors, knee extensors, and knee flexors. We found that there was co-contraction occurring in the TA and the proximal leg muscles (vastus lateralis, vastus medialis, semitendinosis, and biceps femoris), and given the larger volumes of the proximal leg muscles, this likely has a substantial contribution on the whole body energetic cost. Our models do, however, capture the general trends in the data with the effects of muscle activation dominating the changes in energetic cost with both changes in duty cycle and muscle fascicle length.

### 4.2 Mechanics inform energetics

Together, our modelling results demonstrate that the mechanics of muscle contraction is the primary driver of muscle energy use during the range of tasks explored within. Across the three studies, we identified two primary mechanisms that contribute to the observed changes in energetic cost with changes in (i) cycle frequency [van der Zee and Kuo, 2021], (ii) duty cycle [Beck et al., 2020], and (iii) fascicle length [Beck et al., 2022]. In the first two studies, the changes to contraction frequency and duty cycle resulted in an increased strain rate, which, as seen in subsection 3.1 and subsection 3.2, requires higher activation levels to maintain the same level of force. The increased strain rate resulted in greater maintenance heat rates, along with larger relative contributions from the shortening and lengthening heat rates. These increases in shortening heat due to faster shortening velocities, also known as the Fenn effect [Fenn, 1923], have been demonstrated to occur *in vivo* [Ortega et al., 2015]. The second mechanism is the decreased fascicle lengths, which required higher activation levels owing to the leftward shift in the muscle’s operating region on the force-length relationship (see Figure 5). In this case, we found no change in the distribution between the maintenance and shortening heat rates. This is to be expected, as shifts in the fascicle length only resulted in changes to the operating lengths of the muscle and not the strain rates of the fascicles.

### 4.3 Experimental data informing models

In addition to investigating the main drivers behind energetic cost, the other purpose of this study was to (i) investigate the role of both input data on model predictions and (ii) understand *in vivo* experimental data (normalized activation levels, maximum isometric force measures, etc.) needed to obtain accurate predictions of isolated muscle or groups of muscle energetics. We investigated the influence of input heat rate data obtained via isolated fibre bundle experiments from three sources [Barclay, 1996, Barclay et al., 2010, Lichtwark and Barclay, 2010], and illustrated that the range in experimental inputs can have a substantial impact on overall energetic cost. These experiments are highly complex, and only a few select groups engage in such endeavors (for a review see e.g. [Woledge et al., 1985, Barclay and Curtin, 2023]). Hence, caution should be exercised when selecting data to inform model parameters. Further, how energetic rates might vary in human muscle is also unclear. Some studies have investigated heat released in humans *in vivo* [Bolstad and Ersland, 1978], but this can only be done during isometric experiments. They found that for the SOL, the heat rate released during a maximum isometric contraction was *≈* 16 *W*, which is smaller than the *≈* 50 *W* predicted by our model. This could potentially be due to the loss of heat to the environment through the skin, other tissues, or blood flow, which is difficult to limit. Recently, functional phosphorous magnetic resonance spectroscopy (P-MRS) has been validated as a non-invasive measure of muscle-level energy consumption *in vivo* Haeufle et al. [2020]. P-MRS provides a measure of the phosphocreatine breakdown, which is indicative of the ATP hydrolysis (see e.g. Barclay and Curtin [2023]), and thus could provide more reliable *in vivo* data to inform energetic models. The sensitivity analysis of the model (Figure 6) demonstrated not only the role of the energetic parameters on the energetic cost, but the role of the force-velocity relationship parameters. There is a high level of dependence, especially during shortening contractions, as the location of the muscle on the force-velocity curve will drive changes in muscle activation level.

In addition to the intrinsic muscle parameters that can be obtained from isolated fibre-bundle experiments, there are data that could be non-invasively measured during *in vivo* human experiments that could improve the modelling predictions. One key contributing factor is co-contraction, which can drive changes in both the magnitude and the qualitative trends in experimentally measured energetic costs. To limit this, researchers could provide biofeedback on co-contraction levels to the participant or focus on single muscle joints (e.g. abduction of the second digit in the hand is done solely by the first dorsal interosseous). Another variable is the maximum isometric force produced by the muscle. The maximal torque at the joint level needs to be reported as this will alter both the mechanics, through the prediction of muscle activation, and the scaling of the energetic rates. Reporting muscle activation data that is normalized to maximum voluntary contractions also provides a key measure to compare with model predictions. One aspect that was not investigated in this study, but will likely play a role in energetic cost predictions at higher intensities is the role of motor unit recruitment. Given the large difference in energetic rates between slow and fast-type muscle fibres (see Table 2), it is likely that the timing of motor unit recruitment will substantially influence energetic cost [Swinnen et al., 2024].

### 4.4 Limitations and future directions

The model implemented and tested here has limitations that could be improved in future work. First, the experimental data used as input for the models (e.g. maximum isometric force, optimal fascicle length, etc.) was typically based on data averaged across individuals, in some cases from different studies, and so the parameters used may not represent the experimental sample or an individual subject. Further, there can be large variations in energetic rates (Beck et al. [2020] found metabolic cost could vary by 100*W* between participants in some conditions); thus, this could likely be improved upon by predicting energetic cost on a subject-specific basis. Second, this study used a simplified Hill-type model to describe the mechanics of muscle contraction, which fails to account for effects from the architecture of muscle, muscle mass, and detailed tendon mechanics. We found that shortening rate plays an important role in muscle energetics, but we do not take into account the effects of muscle gearing, owing to multi-dimensional shape changes (see eg. Kelp et al. [2023]). To better capture this aspect of muscle contraction, we could implement a three-dimensional model of muscle (eg. Blemker and Delp [2005], Dick and Wakeling [2018], Wakeling et al. [2020]), which would allow us to account for not only the effects of architecture on the muscle fascicle shortening but also the energy required to deform the muscle itself [Wakeling et al., 2020, Ross et al., 2021, Ross and Wakeling, 2021]. Another aspect that has been simplified in the model is the use of a constant tendon stiffness. Tendon stiffness has been shown to play a role in the energetics of muscle [Lichtwark and Wilson, 2007], and a more accurate model that accounts for nonlinear tendon properties could alter the energetic rates predicted by the model.

Future studies could also include a motor unit recruitment model (e.g. Fuglevand et al. [1993], Caillet et al. [2022]) to better capture the energetic cost at higher levels of activation. In the simulations here, the contractions were either at very low activation levels (vdZ2021) or in muscles with very low proportions of fast fibres (B2020 and B2022), thus a recruitment model was not implemented for this study. In addition to the recruitment model, accounting for excitation-activation coupling (eg. Mayfield et al. [2023]) may provide more realistic energetic costs during cyclical contractions, where the Ca^2+^ concentrations may not return to a baseline level over multiple contractions. Models that are more biophysically motivated, which include detailed activation dynamics, have been developed for whole muscle simulations [Ma and Zahalak, 1991], but are not as widely utilized in whole body simulations of movement.

## 5 Conclusion

The physiologically-based muscle model used in this study was able to capture the trends in muscle energy use observed during *in vivo* tasks across a range of contraction conditions [van der Zee and Kuo, 2021, Beck et al., 2020, 2022]. The model was unable to accurately capture the absolute value of the experimental energetic rates, which could be due to many factors, such as co-contraction of other muscles, estimated maximum isometric force values, or motor unit recruitment. To improve modelling approaches, future experiments should endeavour to report accurate measures of these factors, which likely contributes to the overall energetic cost. Our results indicate that these models could be improved upon by including a motor unit recruitment model or a more detailed mechanical model. This study details a modelling framework that utilizes *in vivo* experimental measures to capture muscle energetics and provides rich insights into the mechanics of muscle contraction that models should account for when estimating single muscle energy use.

## 8 Data availability

All relevant data can be found within the article and its supplementary information.

